# The dual Ewald sphere reconstruction for cryoEM

**DOI:** 10.64898/2026.06.24.734255

**Authors:** J Bernard Heymann

## Abstract

Images in the electron microscope are formed by electron scattering and focusing. The spherical geometry of these processes gives rise to two coherent, conjugate spherical wave fronts, known as Ewald spheres. These spheres are associated with the two halves of the contrast transfer function (CTF), and their widths are determined by the focal gradient through the specimen. To properly correct for the CTF, each half of the CTF must be applied to an image individually and integrated into the reconstruction into the corresponding Ewald sphere. Theory indicates that this dual Ewald sphere reconstruction method should recover the maximal amount of information possible. This method was compared to the other reconstruction methods commonly used: the projection approximation (ignoring the Ewald sphere), the simple insertion and the single sideband methods. In simulated reconstructions the dual Ewald sphere method recovered the most information when the correct half of the CTF is matched to the corresponding Ewald sphere. If the wrong half is matched, the result worse than the projection approximation method. Examining reconstructions from real data indicated that the dual Ewald sphere method performs at least as well as the simple insertion method, but not as good as in simulations. The likely reason is the two-fold ambiguity in the assigned orientations of the particle images, which remains an issue to pursue in further studies. In conclusion, the dual Ewald sphere reconstruction method may offer the best way to calculate very high resolution reconstructions when the micrograph quality warrants it.

**Highlights:**

1. The dual Ewald sphere reconstruction corrects for the two halves of the CTF.
2. The signs of the two halves of the CTF must correspond to the focal gradient.
3. Determining the focal gradient for individual particle images remains unresolved.
4. Complex reconstructions indicate any real space phases are artifacts.

## Introduction

The reconstruction of cryoEM maps have reached atomic detail in many studies (examples: (Fromm et al., 2023; Kucukoglu et al., 2024; Savva et al., 2024; Xie et al., 2017)). For most studies a resolution of 2-3 Å is adequate to build atomic models, providing the necessary information for typical biological questions. However, the effort to achieve the highest possible resolution also improves the quality of lower resolution maps. In this we still have several open questions to both improve data processing and better understand the physics behind it. One of these is the proper handling of the Ewald sphere that is a direct consequence of the spherical geometry of electron scattering and focusing (Heymann, 2023; Heymann, 2026). Because of the high acceleration voltages used, the Ewald sphere is relatively flat and can be approximated as a plane, known as the “projection approximation” (equivalent to Fraunhofer diffraction). Therefore, many reconstructions are done by simply integrating the Fourier transform of a particle image into the central section of a reconstruction in the proper orientation.

The pertinent change came with the realization that the Fourier transform of a particle image should be integrated along the dual Ewald spheres, implemented as the “simple insertion” method (Wolf et al., 2006). In this method, the CTF is still corrected in its full form before the integration. An alternative method is to consider the two halves of the CTF as manifestations of scattering to two sides of the image as original suggested by DeRosier (DeRosier, 2000), resulting separable “side bands” (Russo and Henderson, 2018). This method is in widespread use in the Relion package (Zivanov et al., 2018), but it has an inherent discontinuity that creates artifacts (Heymann, 2026).

The Ewald sphere effect is related to the thickness of the specimen (Heymann, 2023). Therefore, in another approach, the idea is to correct for the CTF in the projection approximation by reconstructing smaller blocks of the specimen with assembly afterwards. This “block-based” reconstruction (Zhu et al., 2018) pushes the resolution limit of the projection approximation further out but does not address the nature of the Ewald spheres.

In a previous paper I used the simple insertion method to illustrate that the Ewald sphere does not pose a limit on the resolution of a cryoEM map (Heymann, 2023). In a subsequent paper I examined focal series micrographs to understand how spherical scattering and focusing that result in the conjugate Ewald spheres relate to the CTF (Heymann, 2026). I showed that applying one half of the CTF flattens one Ewald sphere, while doubling the curvature of the other sphere. This suggested a way to correct for the CTF along the Ewald spheres by the two halves of the CTF, which should yield better reconstructions.

In this study I propose the dual Ewald sphere reconstruction method in contrast with the other methods. Importantly, I show that the signs for the two CTF aberration functions must match the focus gradient of the Ewald spheres to recover maximum information. The dual Ewald sphere reconstruction with the correct signs always exceeds the other methods with simulated images, with the simple insertion and single sideband methods equivalent to a random assignment of the signs. Reconstructions from real data is complicated by other inherent resolution-limiting factors, resulting in difficult interpretation. One feature is that the direction of the electron beam with respect to the projection image is two-fold ambiguous. This affects the distinction between the two Ewald spheres and how to appropriately correct them. Efforts to assign the orientation to the correct beam direction in real data was not successful, likely because of inadequate quality in the micrographs. The conclusion is therefore that the dual Ewald sphere reconstruction is at least as good as the simple insertion method, and should be valuable for very high quality micrographs of large specimens.

## Reconstruction theory

### The nature of the signal

Where the electrons hit on the detector in the electron microscope depends on how they are scattered and focused. The scattering geometry generates a spherical wave front, the Ewald sphere, where all the scattered electrons are in phase (Ewald, 1913; Ewald, 1969). Therefore, the hits in a micrograph form a coherent pattern consistent only with the two Ewald spheres associated with scattering and focusing. The widths of these spheres are determined by the focal gradient through the specimen, and thus the specimen thickness, limiting the extent to which the projection approximation can be used (Heymann, 2023). Therefore, the coherent signal on the detector can be represented as a superposition of the ridge peaks of the two Ewald spheres (Heymann, 2026). Given the location {*u*, *v*} in the Fourier transform of the micrograph at spatial frequency 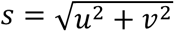, the Ewald spheres are offset from the central section by 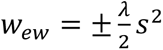 with *λ* the electron wavelength. The coherent signal recorded in the micrograph is effectively the sum of the Ewald sphere ridges modified by the CTF phase shifts ±*γ*(***s***):

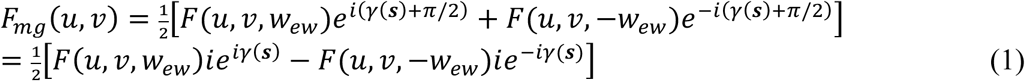

The phase shift is typically modeled with three major terms:

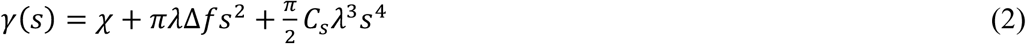

where *χ* is a frequency-independent phase shift term, Δ*f* is the average defocus, and *C_s_* is the spherical aberration. The constant phase shift, *χ*, is the amplitude contrast, where its magnitude is a function of the refractive index of the specimen. Note that *ie^iγ^*^(***s***)^ = *e^i^*^(*γ*(***s***)+*π*/2)^, accounting for the fact that the image is mostly imaginary. Other terms that may contribute to the CTF (such as astigmatism and higher order aberrations) can be added to the phase shift to compensate for further aberrations without changing the principles here.

In the projection approximation, *w_ew_* is assumed to be small enough to set it to zero:

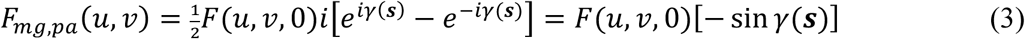

This approximation is valid up to the point where the Ewald spheres separate. The Ewald sphere ridge is modeled as a sinc function, where the width is determined by the focal gradient through the specimen, and thus its thickness (Heymann, 2026). The separation point is defined as the first node where the two sinc functions are zero, where, for a given specimen thickness t, the spatial frequency is (Heymann, 2023):

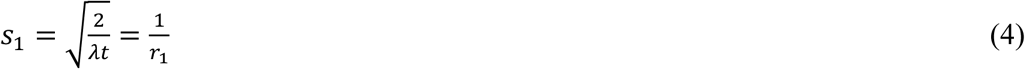

giving a resolution limit, *r*_1_, for the projection approximation. Beyond this point phase reversal occurs that decoheres the signal on the central section in frequency space that limits the resolution of the projection approximation.

The problematic parameter in these equations is the effective thickness of the particle, t. This derives from the distribution of atomic coordinates in the direction of the beam. One way to estimate it is to use the calculation that leads to the sinc function, and then determine the nodes at frequencies *s_n_*:

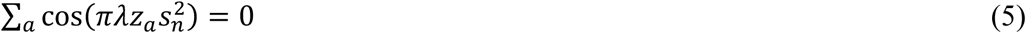

The result is checked for the first point at which it is less than zero to determine *s*_6_. The effective thickness is then calculated based on equation 4.

### CTF correction and the recovery of information

#### CTF correction using the full function

The traditional approach in cryoEM is to add the two halves of the CTF and correct using the sine function as shown in equation 3. This is used both for the projection approximation and simple insertion reconstructions. If we apply the full CTF equation to the model in equation 1, we get:

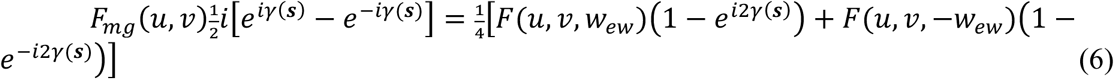

The result is a superposition of the two corrected Ewald spheres as well as doubly imposed phase shifts in opposite directions. This effect was illustrated in the CTF correction of focal series (see (Heymann, 2026) Figure 5d). The phase shifts of ±2*γ*(*s*) generates the so-called “twin image” (Downing and Glaeser, 2008) that was also discovered in the original development of inline holography (Gabor, 1948). Only a quarter of the information is correct for each Ewald sphere, with half being correct where they overlap. This works in reconstruction because only the coherent part contributes and the rest is averaged out. The only difference between the projection approximation and simple insertion methods is that the integration in frequency space is in the central section in the former and along the Ewald spheres in the latter (Heymann, 2023; Wolf et al., 2006).

#### Dual CTF correction using the two halves of the function

Russo and Henderson first considered the two halves or aberration functions of the CTF separately (Russo and Henderson, 2018). In their method they preserve Friedel symmetry throughout, so that the first half is applied to one half of the image, and the conjugate half to the other half of the image (the “single sideband” method). This creates a hard boundary where the sign switches, introducing problems in the calculation. Their solution to do the application in sectors, however creative, does not avoid the hard boundary (Heymann, 2026). Their insistence on maintaining Friedel symmetry is also contrary to the understanding that the individual Ewald spheres do not preserve Friedel symmetry in 2D.

A view that is more consisted with the Ewald sphere as it relates to X-ray crystallography, is to associate the two halves of the CTF with the two Ewald spheres (Heymann, 2023; Heymann, 2026). One half is applied during reconstruction for one Ewald sphere and the conjugate half for the conjugate Ewald sphere. Applying the aberration functions separately gives the following pair of equations:

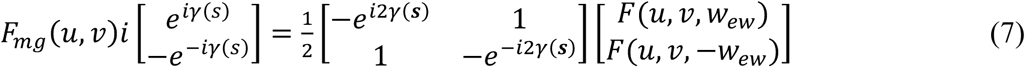

Notice the correspondence of the terms to those in equation 6. For each of the functions, half of the information for each Ewald sphere is recovered, while the other half is subject to double the oscillations of the CTF. The latter is not coherent over the many different instances of particle orientations in reconstruction and effectively contributes to the noise. Unfortunately, the matrix in equation 7 is singular and therefore cannot be inverted.

Applying equation 7 requires that the correct half of the CTF must be matched with the correct Ewald sphere to be consistent with the gradient of focus. In the single sideband method, only half of the image is corrected properly to match the focal gradient, so that it suffers the same loss of information as the simple insertion method (only ¼ is recovered). It should therefore be possible to recover at least half of the information with applying the two halves of the CTF for the two Ewald spheres separately during reconstruction.

## Reconstructions from simulated images

The value of simulated images is in testing code written to implement equation 1 for imaging and assess the various reconstruction methods. It is also the only way to determine the recovery of information by comparing the reconstruction with the original atomic model. For real data where the exact model is not known, the usual test in one of consistency between halfmaps (two reconstructions from mutually exclusive halves the data). Images were generated according to equation 1 with a range of defocus values, and then used to assess the different types of reconstruction:

1. Projection: CTF corrected by the sine function (integrated into the central section).
2. Insertion: CTF corrected by the sine function.
3. Sideband: CTF corrected according to the Russo & Henderson scheme (Russo and Henderson, 2018).
4. Dual: CTF corrected by the individual CTF halves:

a. Correct signs
b. Wrong signs
c. Random signs

Note that except for the projection approximation (method 1), all methods integrate along the Ewald spheres.

### Recovery of information

To simplify the analysis, I first start with images without any noise. The chosen example is the adeno-associated virus (AAV) case with an effective thickness of ∼215 Å for the capsid shell (Xie et al., 2017). The phase flip in the projection approximation occurs at ∼1.9 Å at 100 kV, ∼1.6 Å at 200 kV, and ∼1.4 Å at 300 kV. The best resolution obtained is 1.56 Å at 300 kV (Xie et al., 2020), which still places it before the phase reversal. Thus, a sampling of 0.5 Å/pixel was selected to be able to observe the effects to the Nyquist frequency of 1 Å for images generated at 200 kV. Only 100 randomly oriented noise-less projection images were simulated, because the high symmetry and absence of noise ensure good coverage in frequency space.

Images were generated with a negative setting, i.e., the upper Ewald sphere was multiplied with the aberration function *e*^−*i*(*γ*(*s*)+*π*)^, and the lower with its conjugate. The reconstruction done with a positive setting gave the best recovery of information as assessed by resolution, assigning it as the one with the “correct signs”, while the one done with a negative setting was assigned as the “wrong signs”. This is a sanity test that indicates the calculations give the expected result.

Figure 1 shows the comparative results for the various AAV reconstructions. The biggest difference in the sharpness of features in the maps is between the dual Ewald sphere reconstructions with correct and wrong signs (Figure 1A,B). This is particularly evident in the enlarged views (Figure 1F,G), with the other reconstruction methods falling in between (Figure 1C-E). It is remarkable that at a resolution of ∼3 Å, the incorrect map still resolves individual atoms, just not very well.

**Figure 1:**
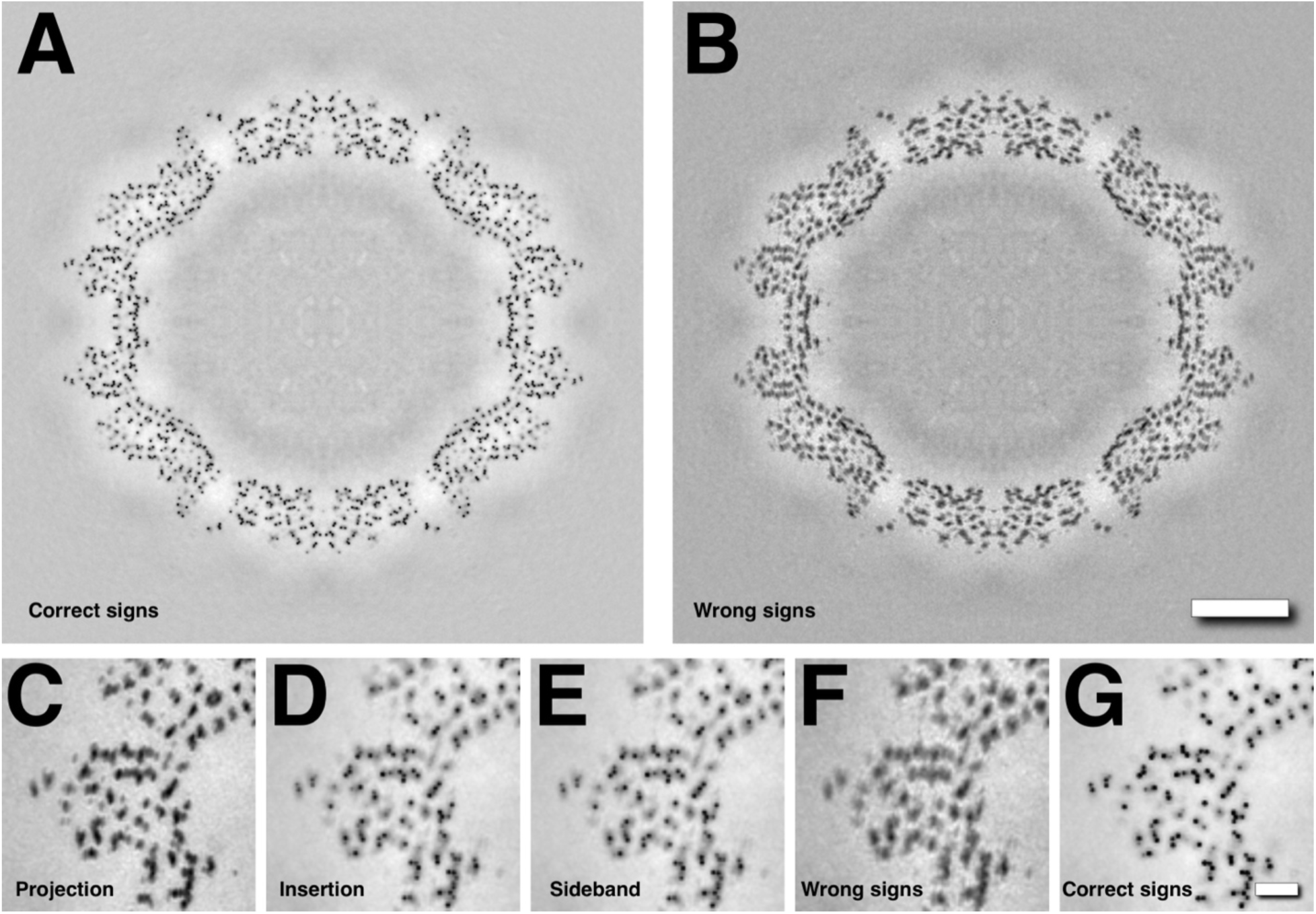
AAV reconstruction with the (A) correctly signed and (B) incorrectly signed application of the two Ewald sphere halves of the CTF. One hundred AAV capsid images were simulated with the program **bess** according to equation 1 at 200 kV at a pixel size of 0.5 Å and with CTF parameters, amplitude contrast 0.07, spherical aberration (Cs) 2.7 mm, and focus range −0.5 – −1.5 µm. Reconstruction was done with the program **breconstruct** to a resolution of 1Å and with icosahedral symmetry imposed. Scale bar: 50 Å. (C-G) Enlarged parts of five reconstructions to compare the appearance of atoms: (C) Projection approximation; (D) Simple insertion method; (E) Single sideband method; (F) Dual Ewald spheres with the wrong signs; (G) Dual Ewald spheres with the correct signs.

While the appearances of these maps are informative, the nuanced differences are better interpreted from statistical comparisons, including Fourier shell correlation (FSC) with the simulated 3D map and the recovery of intensities indicated in the radial power spectrum (RPS) (Figure 2). The reconstruction according to the projection approximation shows the phase reversal (flip) at ∼1.6 Å (Figure 2A, grey curve). As expected, the reconstruction with signs that match those of the simulated images gives a close to perfect map (Figure 2A, blue curve). In contrast, the reconstruction with the wrong signs is worse than the projection reconstruction, with the phase flip at ∼2.3 Å (which is ∼1.6√2Å) (Figure 2A, red curve). Also note that the difference in choosing the signs already has an impact at a much lower resolution of ∼4 Å. Interestingly, the simple insertion (green curve) and single sideband (orange curve) methods give almost identical results, indicating that they integrate the same amount of information. Reconstructing with random signs with half correct comes close to the insertion and sideband methods (Figure 2A, purple curve). In all three the latter cases the theory indicates that only about a quarter of the coherent signal contributes to the reconstruction, while for the dual Ewald sphere reconstruction with the correct signs, about half of the information is recovered. Examining the radial power spectra of the reconstructions (Figure 2B) confirms the relative incorporation of the quantitative information, where the differences start at ∼4 Å. While the insertion and sideband methods give reasonable reconstructions, they always show lack of complete information recovery as can be illustrated by reconstructions from different particle numbers (Figure 2C). At high frequencies, only the dual Ewald sphere reconstruction with the correct signs reaches completion at high particle numbers (Figure 2D). This shows the main benefit from the correct reconstruction is in using the maximal possible information in the particle images. For practical purposes, this means that with the dual Ewald sphere method, several fold fewer particles would be required to reach the same resolution.

**Figure 2:**
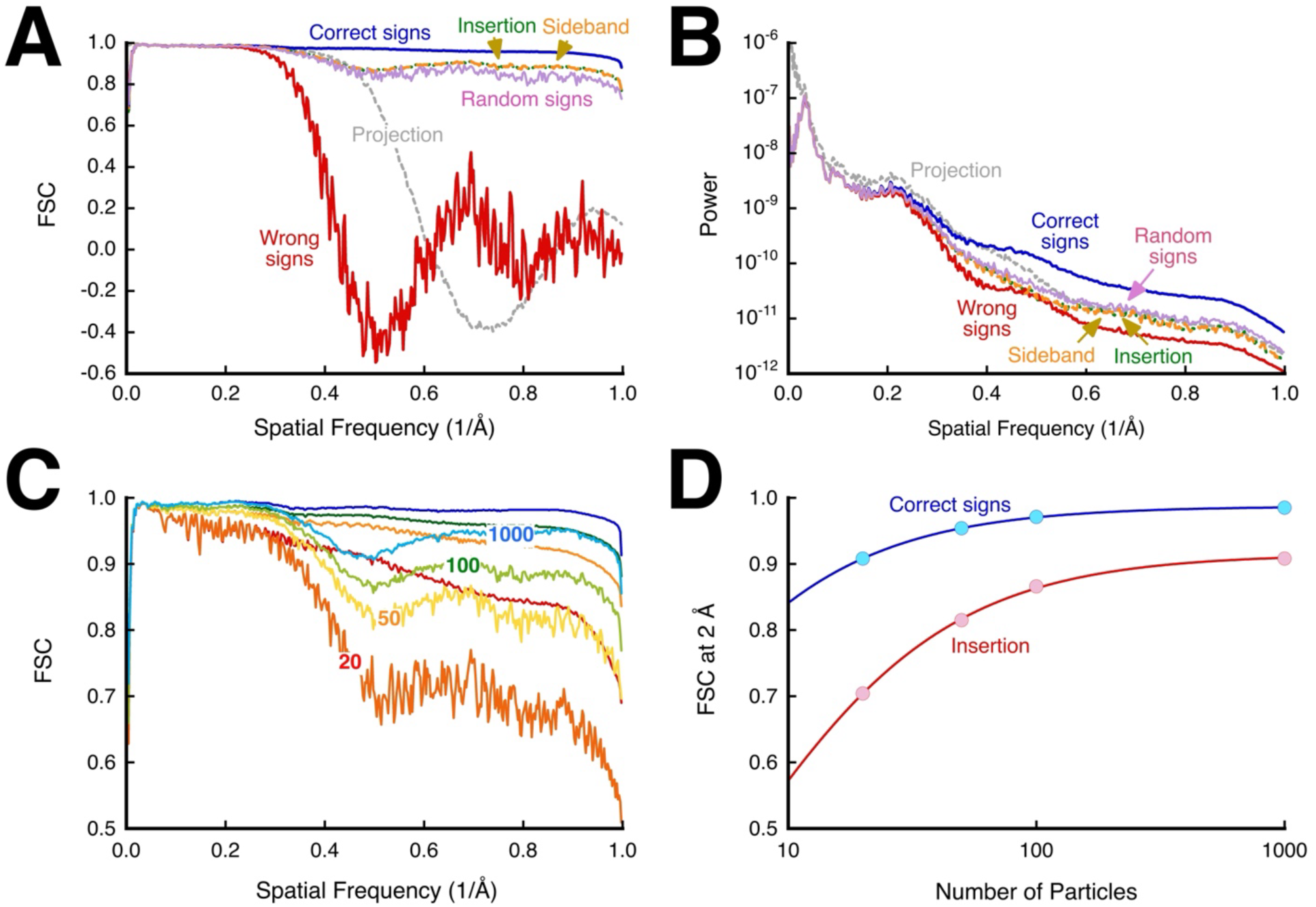
Analysis of reconstructions of simulated images of AAV (as in Figure 2). The Fourier shell correlation (FSC) curves were calculated using the program **bresolve** and the radial power spectrum curves were calculated using the program **bradial**. (A) FSC between the original potential map and the reconstructions calculated according to the projection approximation (gray), the simple insertion method (green), the single sideband method (orange), and dual Ewald spheres method with correctly (blue), incorrectly (red) and randomly (purple) signed spheres. (B) Radial power spectra of the reconstructions to show the quantitative recovery of the signal. The power scaling is arbitrary. (C) Dual Ewald sphere and simple insertion reconstructions from different numbers of particles (20,50,100,100). (D) The FSC at 2 Å from panel C showing the lower recovery of information for the insertion method compared to the dual Ewald sphere method. The fits are modeled as a function of the number of particles, n: 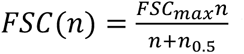, with the parameters for insertion: *FSC_max_* = 0.91 and *n*_0.5_ = 5.99, and for dual Ewald: *FSC_max_* = 0.99 and *n*_0.5_ = 1.74.

### Consistency of information

The usual assessment of cryoEM maps is to compare reconstructions done from two different sets of particle images, typically half of the data and producing the so-called halfmaps. This tests the consistency of the reconstructions and is not the same as recovery of information tested in the previous section. Because there is typically some crosstalk between the processing of half-sets of particle images, this measure often contains some bias. In our case here with simulated images, the orientations for the particle images are known and therefore alignment is not an issue.

Figure 3A shows that the correct and wrong signs of the dual Ewald sphere reconstruction method in our AAV test case gave the best and worst results, respectively. Even the projection approximation is better than choosing the wrong signs for the dual method. The single sideband method is equivalent to choosing the signs randomly, while the simple insertion method fared better. With a larger number of particles, the simple insertion method comes close to full consistency, but still lower than the dual method with correct signs (Figure 3B). The real advantage of the dual method shows with smaller numbers of particles (Figure 3C). The introduction of noise of course limits the resolution, but the trends between the reconstruction methods remain (Figure 3D).

**Figure 3:**
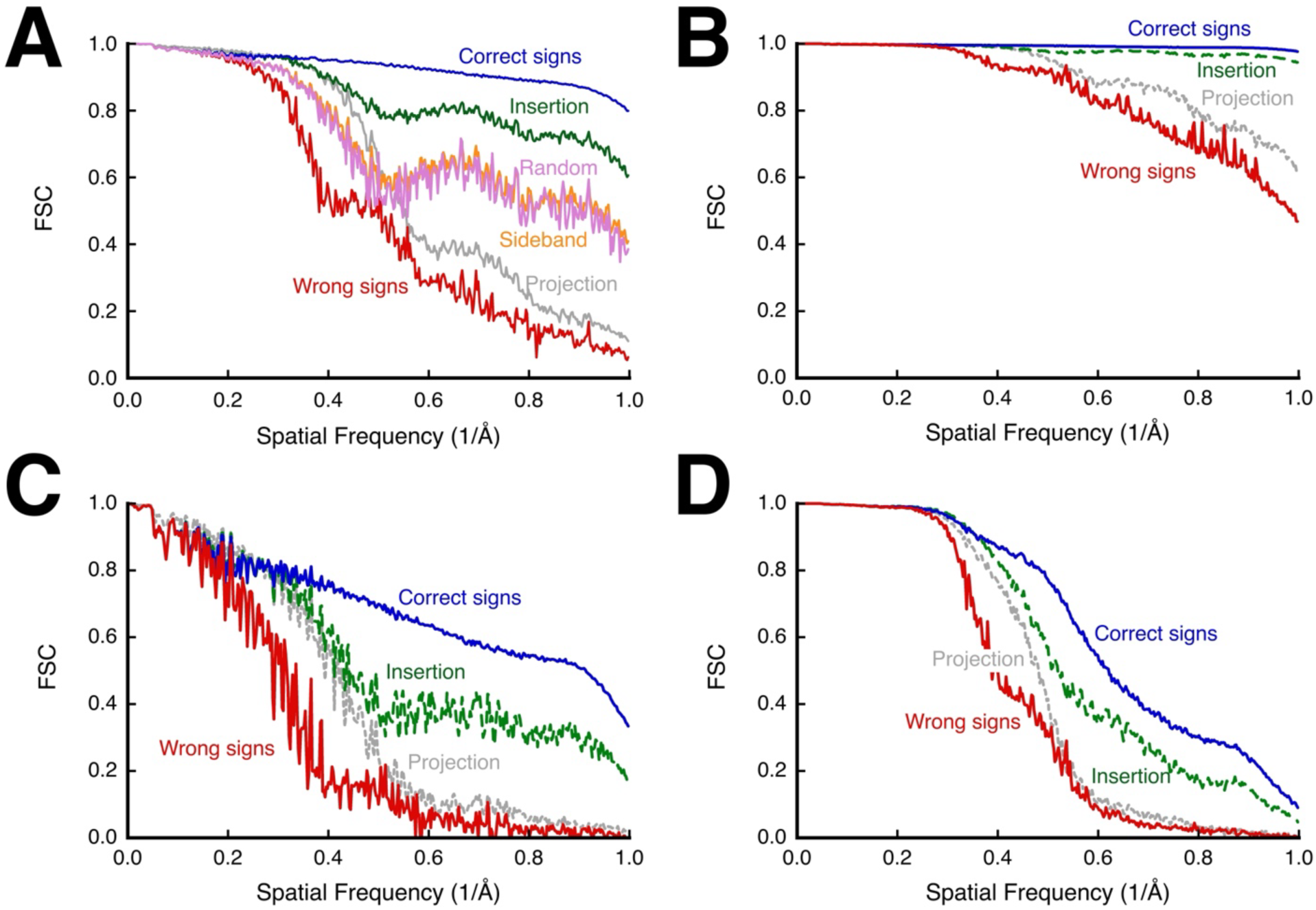
Consistency of information assessed by comparing halfmaps of the AAV simulated images (as in Figure 2). (A) 100 images. (B) 1000 images. (C) 20 images. (D) 1000 images with Poisson distributed noise at an SNR = 0.1.

## Reconstruction from real images

### The caveat of simulated images

The imaging model proposed here is based on an interpretation of the results from a previous simulation study (Heymann, 2023) and reconstructions from focal series of micrographs (Heymann, 2026). Because the images are simulated in a particular way, the recovery of information is some sense guaranteed. In the dual Ewald sphere method, the choice of signs is matched to the choice made during image simulation. However, in a real case, it is unknown which sign choice will produce the best results. From the simulation tests, one would expect a clear distinction whether the correct or wrong signs were chosen.

### The caveat of recovery from real images

Recovery of information in real cases requires an accurate atomic model, but such a model is still based in resolution-limited data. Nevertheless, assessing different reconstruction methods against the same model still provides a relative comparison regarding recovery of information. Such a comparison is expected to be consistent with the correlation between halfmaps as is typically done in real cases.

### A case of a small molecule, ß-galactosidase

An available set of particle images ß-galactosidase with associated processed parameters (Merk et al., 2020) was used to test the different reconstruction methods. The choice of signs for the dual Ewald sphere method that resulted in the best reconstruction was the same for recovery (Figure 4A), and consistency between halfmaps (Figure 4C), even with small numbers of particle images (Figure 4E). All Ewald sphere methods fared better than the projection approximation. Even the dual method with the wrong signs produced a better reconstruction. In this case the simple insertion method coincides with the dual method with the correct signs, while the single sideband method is slightly worse. The conclusion is that the particle is small enough with an effective thickness of 93 Å so that the difference between the Ewald spheres is very small. While phase reversal occurs at ∼1.08 Å, the curve for the projection approximation reconstruction separates from that of the others at ∼2.5 Å for a large number of particles, and at an even lower resolution (∼3 Å) for only 10000 particles (Figure 4E). This means that attending to the Ewald sphere is already beneficial at lower resolutions.

**Figure 4:**
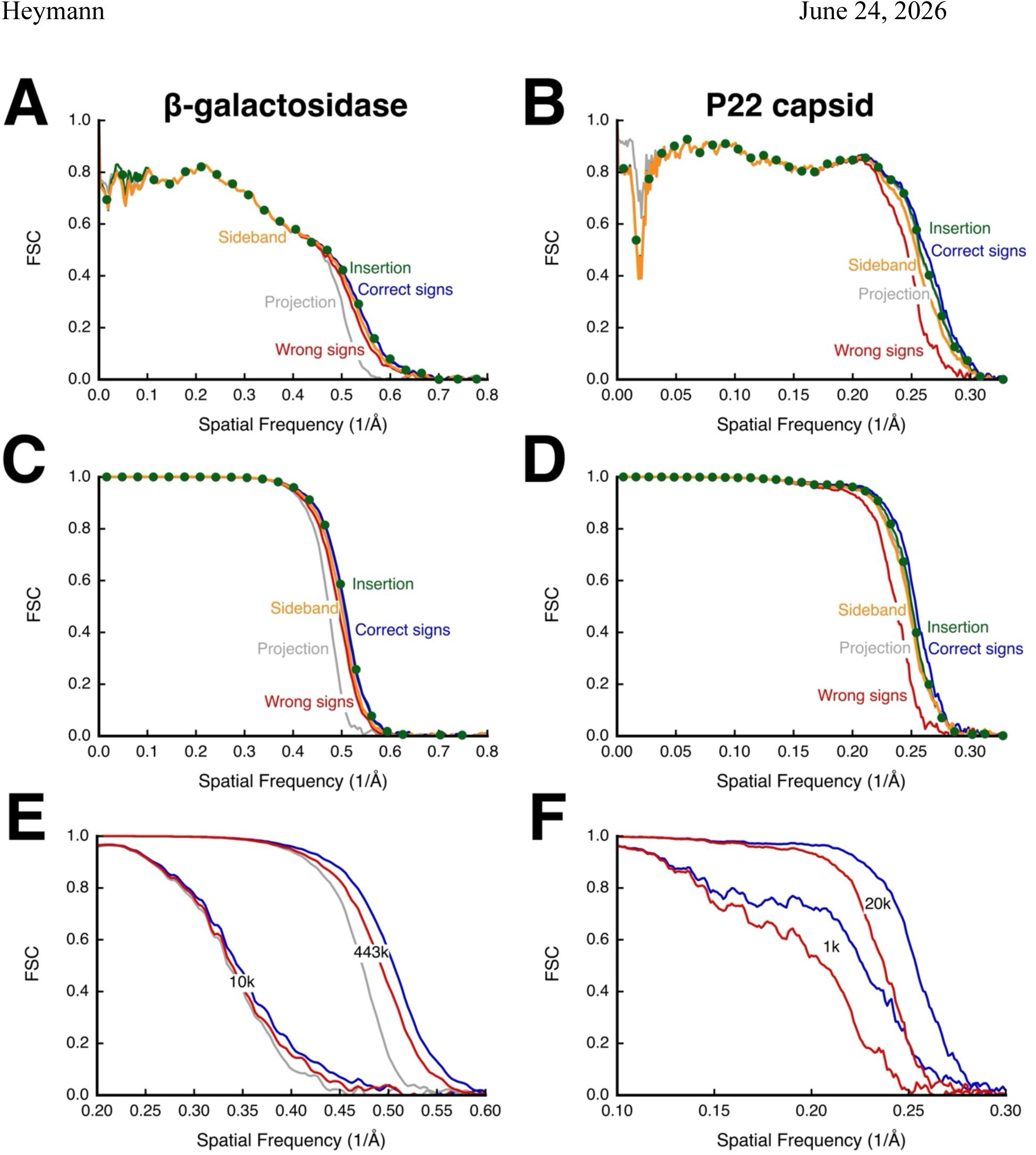
Comparisons of reconstructions of (A,C,E) ß-galactosidase with D2 symmetry from available images (EMPIAR: 10446) (Merk et al., 2020), and of (B,D,F) bacteriophage P22 capsid with icosahedral symmetry from available images (EMPIAR: 10083) (Hryc et al., 2017). (A,B) Recovery of information assessed by FSC against the atomic models (PDB 1JYX and 5UU5, respectively) for five reconstruction methods. (C-F) Consistency of information assessed by FSC between halfmaps. (E,F) Reconstructions from small or large numbers of particles preserve the relative comparison between correct sign (blue) and wrong sign (red) reconstructions (k indicates 1000s). Colors indicate the different reconstruction methods: gray: projection approximation; green (dots): simple insertion; orange: single sideband; blue: correct signs; red: wrong signs.

### A case of a large particle: bacteriophage P22 capsid

Because the Ewald sphere becomes more important for larger particles, I retrieved an available data set of particle images taken at 300 kV of the bacteriophage P22 capsid with an effective thickness of 588 Å (EMPIAR: 10083) (Hryc et al., 2017). While the authors advertised that these images have embedded orientation parameters, it was not possible to recover them. Instead, I aligned the images against a reference map calculated from the atomic coordinates (PDB 5UU5). The reconstructions show the following series of improving resolution (Figure 4B,D): wrong signs dual method, single sideband method, projection approximation = simple insertion, and correct signs dual method. The frequency at which phase reversal occurs is calculated as 2.4 Å. For 20k particles, the differences between the projection approximation and the other methods were already evident at 4 Å. The correct and wrong signs dual Ewald sphere methods separate at 6 Å for 20k particles, and 8Å for 1k particles (Figure 4F). This strongly indicates that the dual Ewald sphere reconstruction is already beneficial for small numbers of large particles at much lower resolutions than expected. It also confirms that the model of image formation implied by equation 1 provide a better correction of the CTF compared to the other methods.

## Testing the ambiguity in projection direction

The choice of signs for the dual Ewald sphere method is applied to each particle image with the assumption that the focal gradient is the same in all micrographs. However, reconstructions from the real data show a much smaller distinction between the “positive” and “negative” applications of the CTF halves than expected from the synthetic reconstructions. One possible reason is that the orientation assignment of the particle images has an inherent ambiguity, in that the direction of the beam (up or down) is unknown. This determines which of the CTF halves should be considered for which Ewald sphere. I explored many variations in trying to classify particle images as “up” or “down” images, based on comparison with the Ewald spheres in a reference map. One promising test is based on modifying a particle image with each half of the CTF, and correlate the Fourier transforms of the two resultant images with the corresponding Ewald spheres extracted from the reference map. Only the parts where the Ewald separate were used in the correlation. The expectation was that if a reconstruction was calculated with particles indicated as “up” or “down” with respect to CTF correction, it would produce a greater distinction between the correct choice versus the opposite choice in the FSC curve. Conversely, if the classification just assigned random “up” and “down” designations, the reconstructions would converge and be similar to the simple insertion reconstruction. This classification indicated a switch of 49% in the ß-galactosidase case, and 25% in the P22 case. Curiously, the FSC curves for the new reconstructions were very close to the original correct and wrong sign reconstructions. Thus, the classification neither selected the correct beam direction consistently, nor did it completely randomize the direction. Two possible reasons are that the required information is masked by noise, or that the classification test is invalid. The former may render it impossible to determine the correct beam direction, while the latter would require further investment in understanding the distinction between the two Ewald spheres.

## Complex reconstruction and real space phases

While the usual reconstruction is done in frequency space and Fourier transformed back to yield a simple real space map, the back transform can be modified to obtain a complex reconstruction. This provides an opportunity to examine the real space phases. Complex maps were calculated using the dual Ewald sphere method for both simulated and real data (Figure 5). These were analyzed by extracting a resolution shell and back transforming to get the real space phases associated with each shell. The rationale is that this separates the real structural information at low frequencies from any non-structural effects at high frequencies. It was found that for simulated maps, the real space phase distribution has an apparent probability of *P*(*ϕ*) = *C*[*ϕ*^−2^ + (*ϕ* ± *π*)^−2^], with the difference with respect to frequency only in the scale *C*. This suggests that this functional form is a consequence of interpolation inaccuracies in the digital Fourier transform itself. This has been confirmed with several maps and sets the baseline for analyzing maps. The reason for the scale difference in the case of the simulated map is likely that only a limited number of images (1000) were used in the reconstruction and the averaging of the real space phases at high frequences was inadequate.

**Figure 5:**
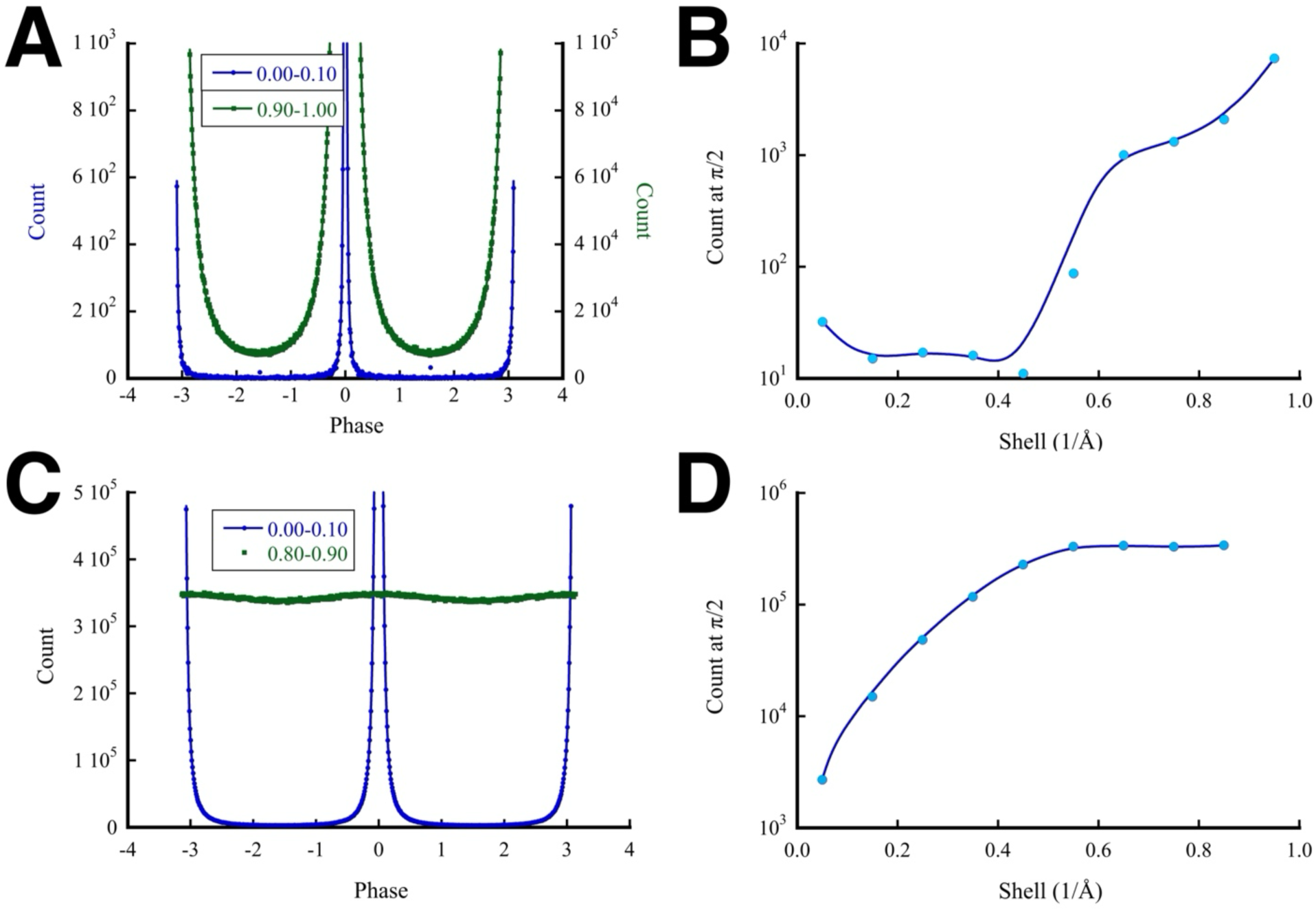
Real space phase distributions for complex reconstructions: (A) A reconstruction of AAV with the correct signs from simulated images, showing the phase distributions for the 0-0.1/Å (blue) and 0.9-1.0/Å (green) shells. (B) The counts of phases for the AAV reconstruction at π/2 (90°) increase in shells where there is a lack of information owing to the small number of particle images used. (C) The ß-galactosidase reconstruction with the correct signs from real images, showing phase distributions for the 0-0.1Å (blue) and 0.8-0.9Å (green). (D) The counts of phases for the ß-galactosidase reconstruction at π/2 increases as the noise increases at higher frequencies. The fits in panels A and C are to: *Counts*(*ϕ*) = *C*[*ϕ*^−2^ + (*ϕ* ± *π*)^−2^] with the scale *C*, 1.186 (blue) and 7829 (green) for the AAV case, and 2680 (blue) for the ß-galactosidase case. The high frequency part (0.8-0.9Å) for the ß-galactosidase case has an almost constant distribution with an average of 3.4 × 10^5^ consistent with it being mainly noise.

Analysis of complex reconstructions from real data clearly show a distinction between the real space phases from low and high resolution shells. Where structural information is clear and well-defined in low resolution shells, the distribution follow that of the simulated maps. At high frequencies, the almost uniform distribution of real space phases is consistent with noise. The conclusion is that the real space phases are very sensitive to noise and lack of data, so that excessive noise and inhomogeneous coverage of the data in frequency space results in real space phases significantly different from zero and π. Therefore, the correct information in reconstructions is real and any residual phases are indications of inaccuracies or noise.

## Discussion

### Dealing with the Ewald sphere in cryoEM

The first important development in handling the Ewald sphere in cryoEM was the recognition that the particle image transform should be integrated into the Ewald spheres rather than the central section (Wolf et al., 2006). This simple insertion method was shown to avoid the resolution-limiting implication in the projection approximation (Heymann, 2023).

The second development was understanding the relevance of the Ewald spheres observed in the 3D Fourier transforms of focal series. I showed that correction for one half of the CTF flattens one of the Ewald spheres in these transforms, while doubling the curvature of the other (Heymann, 2026). This strongly suggested the image formation model expressed in equation 1 and how to correct for that during reconstruction, equation 7.

The third development is presented here in the realization of the dual Ewald sphere reconstruction with correction of the CTF according to equation 7. The choice of which half of the CTF (denoted by the sign of the phase) to apply to which Ewald sphere is important to extract the correct information. In simulations with exact data and parameters, this presents a clear distinction between the different reconstruction methods (Figures 1-3). The dual Ewald sphere method is the only one that can recover the maximal possible information. In real data, several other influences need to be considered, such as the quality of the micrographs and the accuracy of particle alignment. Nevertheless, with the P22 capsid case, the distinction between the correct and wrong sign dual Ewald sphere reconstructions is clear (Figure 4F). This is true even though the resolution is much less than the phase reversal frequency and makes a strong case for the correctness of equation 1.

A potential fourth development to complete the dual Ewald sphere reconstruction method is to reconcile the electron beam direction with the estimated particle orientation. In the real cases analyzed here, the distinction between the correct and wrong signs are not as strong as for the simulated cases. This is also important because the Ewald spheres carry some handedness information. Attempts here to classify the particles into two groups representing the correct and wrong signs indicated that the ambiguity is strong in the ß-galactosidase case and weak for the P22 capsid case. This agrees with the sign distinction results in Figure 4. Therefore, it remains an issue requiring a solution. The problem is that for real images one needs to acquire data that potentially surpasses the frequency at which the phase reverses in the projection approximation. This is difficult because reaching beyond this frequency requires micrograph quality that is hard to obtain.

### Complex reconstruction and real space phases

The images produced in the electron microscope are inherently real, which means that their Fourier transforms have Friedel symmetry. The Ewald spheres map onto the 3D volume, where their conjugate relationship means that Friedel symmetry is retained in the 3D context (Heymann, 2023). Correcting using the two halves of the CTF as described here preserves this conjugate nature. The 3D reconstructions should therefore have minimal real space phases related to the structural information (Figure 5). Deviations in these phases are therefore attributed to noise and computational artifacts. Thus, complex reconstructions may serve as checks on residual noise and problems in reconstruction calculations.

In a recent paper, a method was proposed to determine the handedness of a cryoEM reconstruction from the Ewald spheres effectively using complex reconstructions (Bromberg et al., 2025). Because most mathematical details were not provided, it is difficult to assess the correctness of the method. It is particularly unclear in what form the CTF correction is done. It may be similar to what is done here, but their FSC curves show an inversion in contrast for the wrong matches. This is puzzling, because both the correct and wrong handed maps should show some positive correlation at low frequencies. In the light of the results here that show real space phases in reconstructions are artifacts or noise, it is doubtful that complex reconstructions is useful for either establishing handedness (which can be resolved at low resolution) or handling Ewald sphere issues.

## Conclusion

The dual Ewald sphere reconstruction is based on the interpretation that an electron micrograph is the result of the superposition of the two Ewald spheres associated with electron scattering and focusing. Simulation show that choosing the correct signs for the two halves of the CTF during reconstruction is readily distinguishable from choosing the wrong signs. Reconstructions from real data show some distinction between these choices, suggesting that the image formation model is valid. Nevertheless, there is the potential to further improve on the method if the orientations assigned to the individual particle images can be matched to the focal gradient direction.

## Methods

### Computational details

All computational code was written and executed within the Bsoft package (Heymann, 2018; Heymann, 2022). Projection images were simulated with the programs **bess** and **bsim** (Heymann, 2023), modified to accommodate the application of the two halves of CTF as shown in equation 1. The refinement of aberration parameters for different optical groups (Zivanov et al., 2020) was added to the program **bpart**. The program **breconstruct** was modified to allow the integration of the Ewald spheres with aberration function correction, either with the correct signs, or the switched (incorrect) signs, as well as the simple insertion method (Wolf et al., 2006) and the single sideband method (Russo and Henderson, 2018) or with random selection of the two aberration functions. The reconstructed maps were compared by Fourier shell correlation using the program **bresolve**, and their radial power spectra with the program **bradial**. The reference map was calculated from the atomic coordinates using the program **bpot**.

### Data used

#### ß-galactosidase

The particle stack (442859 images) processed by Merk et al. (Merk et al., 2020) (EMPIAR-10446, **EMD-21995**) was kindly provided by the authors. The original map reconstructed with symmetry D2 in Relion 3.1 gave a resolution of 1.8 Å (58 % of the particles). The micrographs were imaged on a CryoARM200 (JEOL) at 200 kV energy filtered with a slit width of 30 eV and recorded on a K3 camera (Gatan). A range of defocus values from −0.4 to −3.5 µm was used. The original micrographs were nominally sampled at 0.26 Å/px, but the particles were extracted and binned to give a final calibrated sampling of 0.519 Å/px. The particle images underwent polishing in Relion 3.1 (Zivanov et al., 2018) and refinement of CTF parameters for different optical groups with the Bsoft program **bpart** (based on the method of Zivanov et al. (Zivanov et al., 2020)). These were then used to do the various reconstructions to a resolution limit of 1 Å with the program **breconstruct**. Resolution was estimated using the program **bresolve** with a tight real space variance mask generated with the program **bmask** and softened by low-pass filtering to 10 Å with the program **bfilter**. Before refinement of the CTF parameters, the resolution (FSC_0.143_) was recorded as 2.06 Å, while after refinement it was 1.88 Å, and with simple insertion or correct signs dual Ewald reconstruction it was 1.82 Å. The final best reconstruction is therefore close to the original.

#### Bacteriophage P22 capsid

The particle stacks (57292 images) used in the P22 capsid reconstruction (Hryc et al., 2017) were obtained from EMPIAR (10083). The original map was reconstruction with icosahedral symmetry in JSPR to a resolution of 3.3 Å. While the CTF and orientation parameters were supposed to be encoded in the image headers, extracting them with confidence proved to be difficult. I therefore determined the CTF parameters with the program **bctf**, and the particle orientations with the programs **borient** and **brefine**. The reconstructions with the different methods were done with the program **breconstruct**, giving a final resolution to 3.6 Å (FSC_0.143_) for the best map. The difference in the original and final resolutions could be ascribed to parameters used in determining the resolution, such as the mask used and particulars of angular refinement. Because the different reconstructions show consistent patterns relative to each other, I regard the results as representative of the reconstruction methods.

**Table 1:** Data processing information.

| | β-galactosidase | P22 |
| --- | --- | --- |
| EMPIAR code | 10446 | 10083 |
| EMDB code | 21995 | 8606 |
| RCSB code | 1JYX | 5UU5 |
| Number of micrographs | 9134 | 2729 |
| Micrograph size | 11520 x 8184 | 5120 x 3840 |
| Frames per micrograph | 40 | 24 |
| Total dose ($\text{e}^-/\text{\AA}^2$) | 40 | 38 |
| Defocus range ($\mu\text{m}$) | -0.36 – -3.5 | -0.15 – -3.5 |
| Micrograph pixel size ($\text{\AA}$) | 0.26 | 1.07 |
| Particle pixel size ($\text{\AA}$) | 0.52 | 1.07 |
| Particles picked | 442859 | 57292 |
| Box size | 600 x 600 | 864 x 864 |
| Point group symmetry | D2 | I |
| Original resolution<sup>†</sup> ($\text{\AA}$) | 1.8 | 3.3 |
| Resolution<sup>†</sup> ($\text{\AA}$) | 1.8 | 3.6 |

<sup>†</sup> Resolution estimated at $\text{FSC} = 0.143$

## Abbreviations

cryoEM: cryo-electron microscopy
CTF: Contrast transfer function
AAV: Adeno-associated virus
FSC: Fourier shell correlation
RPS: Radial power spectrum

## Acknowledgements

This work used the computational resources of the NIH HPC Biowulf cluster (http://hpc.nih.gov).

